# Defects in the DNA Damage Response of Patient-derived Endometriosis Stromal Cells

**DOI:** 10.1101/2025.10.29.685406

**Authors:** Kora Cadle, Gwen Thomas, Hunter E. Schweiger, Julien Menendez, Rut Molinuevo, Lindsay Hinck

**Affiliations:** Department of Molecular, Cell and Developmental Biology, University of California Santa Cruz, Santa Cruz, California, 95064, United States; Institute for the Biology of Stem Cells, University of California Santa Cruz, Santa Cruz, California, 95064, United States; Genomics Institute, University of California Santa Cruz, Santa Cruz, California, 95064, United States; Laboratory of Molecular Neurobiology, Center for Biomedical Research of La Rioja, Logroño, Spain

## Abstract

With each menstrual cycle, endometrial cells rapidly proliferate and decidualize in preparation for pregnancy. Such rapid proliferation generates replication stress and results in DNA damage with irreparable cells undergoing senescence. Here, we examine the DNA damage response (DDR) of patient-derived stromal cell lines from menstrual effluent (MenSC) of healthy donors and donors with endometriosis. We found that proliferating MenSCs from endometriosis patients (Endo) have a defective DDR that is also present when these cells reach confluence. In G1, these cells contain more 53BP1-nuclear bodies (NBs) and are less senescent than healthy samples. We also treated with hydroxyurea (Hu) to generate replication stress and found that Endo MenSCs responded to this treatment by activating the DDR and generating more 53BP1-NBs. We examined the MRN complex, upstream of the ATM-dependent DDR. Hu treatment of our cell lines resulted in downregulation of all genes encoding the MRN complex, and RAD50 and NBS1 proteins. In a scRNA-seq dataset of endometriosis stromal tissue, we also identified downregulation of *RAD50* and *NBS1*. To evaluate the growth potential of MenSCs, we decidualized cells after Hu treatment and then replated them in growing medium. Untreated endometriosis MenSCs formed more colonies than healthy MenSCs; neither sample type formed colonies after Hu treatment. Together, our studies suggest that endometriosis MenSCs have a defective DDR that may be exploited therapeutically.

## Introduction

Endometriosis is a chronic and debilitating disease characterized by endometrial-like tissue (both glands and stroma) growing outside the uterine cavity; it affects 5-10% of women of reproductive age worldwide (Houston, 1984, Shafrir et al., 2018). The symptoms of endometriosis include chronic pelvic pain and, in many cases, infertility. Despite its high incidence and the negative consequences for patients’ quality of life, it is very difficult to diagnose and there is no cure (Zondervan et al., 2020). Furthermore, the etiology of the disease is uncertain and there may be more than one explanation for its origin (Wang et al., 2020). One widely accepted hypothesis for how endometrial cells populate different regions of the body is Sampson’s theory of retrograde menstruation (Sampson, 1927). This theory postulates that endometrial cells escape the uterus through the fallopian tubes during menstruation, attaching to and proliferating on tissues within the peritoneal cavity. However, it is estimated that 76%-90% of women experience retrograde menstruation, yet far fewer suffer from endometriosis (Halme et al., 1984, Liu and Hitchcock, 1986). This suggests that endometrial cells from women with endometriosis have intrinsic characteristics that allow them to survive, attach, and proliferate outside the uterus.

The endometrium regenerates more than 400 times throughout a woman’s reproductive years. Tissues that self-renew in such a fashion undergo rapid expansion causing cell cycle stress and DNA damage. The human endometrium encounters various sources of DNA damage, including replication stress which has been linked to endometriosis (Munshi and Sachdeva, 2023). Immediately after menstruation, during the initial phase of the regeneration cycle, endometrial stromal cells reline the uterine wall by proliferating rapidly in response to upregulated estradiol. Rapid cell division generates replication stress by depleting replication factors and nucleotide pools, which leads to disrupted replication fork progression and DNA damage. High levels of estrogen are followed by an increase in progesterone that triggers the decidualization of stromal cells to prepare the endometrium for potential embryo implantation. Near the window of implantation, an acute inflammatory stress response occurs in stromal cells that initiates chromatin remodeling and the expression of pro-decidualizing gene networks (Muter et al., 2021, Vrljicak et al., 2018). As these stromal cells decidualize into specialized secretory cells, this reprogramming exacerbates DNA damage from replication stress, resulting in a subpopulation of decidual cells that undergo senescence (Brighton et al., 2017, Rawlings et al., 2021). These pro-inflammatory, senescent cells release bioactive factors, leading to the senescence associated secretory phenotype (SASP) that, along with the withdrawal of hormones, promotes an irreversible inflammatory phase prior to menstruation (Evans and Salamonsen, 2014). However, the nature and extent of the DNA damage response (DDR) in stromal cells of the endometrium has remained elusive (Munshi and Sachdeva, 2023). Studies report conflicting results for the DDR in endometriosis, likely due to differences in sample collection; nevertheless, all studies conclude that the response is altered (Bane et al., 2021, Choi et al., 2018, Hapangama et al., 2008). Thus, the molecular mechanisms governing the DDR in healthy samples and how the response is dysregulated in endometriosis have not been elucidated.

The DDR consists of a coordinated set of signaling pathways that detect DNA damage, signal its presence, and promote its repair or, if damage is irreparable, trigger senescence or apoptosis (Yates et al., 2025). Double-stranded DNA breaks (DSB) are the most detrimental. They are recognized by the MRN complex (MRE11-RAD50-NBS1) that binds the two DNA ends and tethers them together (Syed and Tainer, 2018). MRN facilitates the recruitment of additional repair proteins to broken DNA, notably the DNA repair kinase ATM, leading to its activation (pATM). Activated ATM phosphorylates histone H2A.X at Serine-139 (γH2AX) and the kinase CHK2 that, in turn, activates a variety of checkpoints depending on cell cycle phase. Replication stress, however, stalls replication forks, leading to DNA damage and under-replicated DNA that elude cell cycle checkpoints and persist into mitosis. Should mitotic DNA synthesis fail to complete replication, these DNA lesions are inherited by daughter cells where they are coated in Tumor Protein p53 Binding Protein (53BP1) in G1 of the cell cycle, creating protective nuclear bodies (NBs) (Harrigan et al., 2011, Lukas et al., 2011). Successful repair of these regions with the restart of replication in the successive cell cycle resolves the 53BP1-NB and the DNA damage (Spies et al., 2019). Defective DNA repair, however, can lead to persistent 53BP1-NBs that are observed in aging, senescence, and diseases characterized by a defective DDR (BRCA1/2 deficiency) (Feng and Jasin, 2017, Fernandez et al., 2025, Rasmussen et al., 2016). Still unknown is whether 53BP1-NBs are in endometriotic cells.

In the uterus, tissue resident stem cells are responsible for driving the cyclical growth of the endometrium. These cells reside primarily in the endometrial stratum basalis layer, which is not shed during menstruation. There also exists a population of perivascular, clonogenic stromal cells that reside in the endometrial stratum functionalis. They give rise to stromal cells, which constitute the largest portion of the uterine lining, and are a primary component of endometriotic lesions (Fonseca et al., 2023, Shih et al., 2022). These stromal cells are phenotypically, morphologically, and functionally different in patients with endometriosis (Retis-Resendiz et al., 2025), but the molecular mechanisms by which these differences promote ectopic growth of endometrial tissue in extrauterine environments have not yet been elucidated. It is possible to study these clonogenic stromal cells because they are released in menstrual effluent (MenSCs) and can be cultured *in vitro* (Gargett et al., 2016, Sanchez-Mata and Gonzalez-Munoz, 2021). Given the ease of MenSCs collection and the high likelihood that menstrual fluid’s cellular components directly seed endometriotic lesions (D’Hooghe et al., 1994, Liu et al., 2025, Suda et al., 2018), the study of these cells has the potential to lead to insights into the origin, development, and maintenance of the disease, discovery of potential diagnostic biomarkers, and identification of therapeutic targets. In this study, we investigate differences in the DDR of healthy and endometriosis patient-derived MenSCs, which are thought to have the capacity to undergo retrograde menstruation and initiate endometriotic lesions.

## Methods and Materials

### Collection of human endometrial samples

Menstrual effluent was collected from women with and without endometriosis aged 18 to 45 who were not using hormonal birth control and who provided written informed consent through a UCSC IRB approved study (HS-FY2022-310). Further descriptions of study participants are described in Table 1. Donors used a menstrual cup during night 2 of their menstrual cycle and returned their sample within 24 hours in medium containing DMEM (Thermo Fisher, USA, 11995073) supplemented with 10% FBS (Corning, Mexico, 35-010-CV), 1× antibiotic-antimycotic (ThermoFisher, USA, 15240062), and Fungin (InvivoGen, France, ant-fn-1; 1:1000).

### Endometrial stromal cell isolation

Endometrial stromal cells were isolated as previously described (Sun et al., 2019) with some minor modifications. Briefly, the menstrual effluent was washed with wash buffer [DPBS (Thermo Fisher, USA, 14190-250) containing 5% FBS], pelleted by centrifuging at 300 × *g* for 5 minutes at room temperature, and resuspended in 10 mL of warm 1× trypsin (Gibco, USA, 15090046) to break up the tissue. The cells were washed with wash buffer and centrifuged. The pellet was resuspended in 10 mL of ACK Lysis Buffer (Thermo Fisher, USA, A1049201) and incubated for 5 minutes at room temperature. The suspension was washed with wash buffer and centrifuged. The lysis process was repeated 1-3 times until the pellet was free of red blood cells. Then, the cells were resuspended in medium containing DMEM with 10% FBS and 1× antibiotic-antimycotic, plated in 60 mm dishes, and incubated at 37°C with 5% CO_2_. The next day, non-adherent cells were washed away with DPBS leaving only plastic-adherent stromal cells. The medium was refreshed every 2-3 days until the passage 0 cultures reached 80% confluency. These cultures were expanded to 100 mm dishes using 0.05% trypsin (Gibco, USA, 25300-062). Passage 1 stocks of these cell lines were created for each donor and cryopreserved in liquid nitrogen. A biologically independent sample is defined as the cell line generated from each individual donor. *n*=4 biologically independent cell lines were generated from healthy control samples and *n*=3 biologically independent cell lines were generated from endometriosis patient samples.

### Cell culture

Endometrial stromal cell lines were plated at approximately 30% confluency in growing medium [phenol red-free DMEM/F12 (ThermoFisher, USA, 11039047), 10% heat-inactivated, charcoal-stripped FBS (Biowest, USA, S162C), 1× antibiotic-antimycotic (Thermo Fisher, USA, 15240062), 1× Insulin-Transferrin-Selenium (ThermoFisher, USA, 41400045), 1× GlutaMAX (ThermoFisher, USA, 35050061), 1 nM 17β-Estrogen (Sigma, China, E8875; E2)]. At approximately 80% confluency, the medium was refreshed, and cells were either untreated or treated with hydroxyurea (Sigma, India, H8627), 100 nM Ku-60019 ATMi (Caymen, USA, 17502), 25 µM Mirin (Sigma, USA, M9948) or corresponding DMSO (ThermoFisher, USA, J66650.AD) control. For the hydroxyurea titration, various concentrations (0.1, 0.3, 1, 10, 30 mM) were used. 10 mM Hu treatment was used for all other assays. The cells were collected 48 hours later when they reached 100% confluence or changed to decidualization medium [DMEM/F12, 2% heat-inactivated, charcoal-stripped FBS, 1× antibiotic-antimycotic, 1× Insulin-Transferrin-Selenium, 1× GlutaMAX, 0.5 mM 8-Bromoadenosine 3′,5′-cyclic monophosphate sodium salt (Millipore-Sigma, USA, B7880), and 1 µM Medroxyprogesterone 17-acetate (Cayman, USA, 23664)]. Decidualization medium was refreshed every other day until decidualization day 4. Cells were collected at the indicated timepoints using warm 0.25% Trypsin-EDTA (Corning, USA, 25-053-CI) for downstream analyses. Passage 4-10 cells were used for experiments. Cell lines were routinely checked for mycoplasma (ABM, Canada, Mycoplasma PCR kit, G238).

### Flow cytometry

For EdU incorporation, the Click-iT Plus Flow Cytometry Assay kit (Invitrogen, USA, C10637) was followed according to the manufacturer’s protocol. EdU or corresponding DMSO control was added to the cell medium of proliferating cells (80% confluent) at a final concentration of 10 µM and incubated for 2 hours at 37°C with 5% CO_2_. Cells were collected and the Click-iT reaction was performed according to the established protocol. For MenSC characterization, cultured cells were collected, washed with DPBS, centrifuged at 300 × *g* for 5 minutes at 4°C, and the supernatant was aspirated. Cells were resuspended to a final concentration of 10^6^ cells/mL in DPBS and incubated on ice for 30 minutes in darkness with viability dye (Tonbo, USA, 13-0865-T100; 0.5 µl/mL). Cells were washed with wash buffer (DPBS containing 5% FBS) and centrifuged at 300 × *g* for 5 minutes at 4°C. The cells were incubated in 100 µL of wash buffer with anti-CD140b (Biolegend, USA, 323608; 1:200), anti-SUSD2 (Biolegend, USA, 327406; 1:200), anti-CD146 (Biolegend, USA, 361003; 1:400) on ice for 30 minutes in darkness. The cells were washed with wash buffer, centrifuged, fixed in 1% PFA (ThermoFisher, USA, J61899.AK), and stored in darkness at 4°C until analysis. For cell cycle analysis, cultured cells were collected, washed with DPBS, centrifuged at 300 × *g* for 5 minutes at 4°C, and the supernatant was aspirated. Cells were immediately fixed by vortexing vigorously and adding ice-cold 70% ethanol dropwise for 1 minute. The cells were washed with 1× PBS to remove the fixative and centrifuged. The cell pellet was resuspended in propidium iodide solution [1× PBS containing 25 µg/mL of propidium iodide (Invitrogen, USA, P3566) and 50 µg/mL of RNAse (Invitrogen, USA, 12091-021)] and incubated overnight in darkness at 4°C. Before flow cytometry analyses, cell suspensions were filtered through a 70 µm cell strainer and then analyzed using a Cytoflex LX (6-L UV) cytometer. Populations were analyzed using FlowJo. For all analyses, MenSCs were selected using SSC-A versus FSC-A and doublets were removed by SSC-A versus SSC-H (Suppl Figure 1A). Gating for MenSC characterization used a fluorescence minus-one (FMO) control as the negative population (Suppl Figures 1B, 1D, 1F). For cell cycle analysis, dead cells and doublets were further removed using PI-W versus PI-A and phases were gated based off of the PI-A histogram (Suppl Figure 1G). Gating for EdU positivity used a DMSO control as the negative population (Suppl Figure 1J).

### MTT assay

The MTT cell viability assay kit (Biotium, USA, 30006) was followed according to the manufacturer’s protocol. Cultured cells were plated in a 96-well plate and treated in duplicates with various concentrations (0.1, 0.3, 1, 10, 30 mM) of hydroxyurea (Sigma, India, H8627) or water control when they reached 80% confluency. After 48 hours of treatment when the cells reached 100% confluence, the cells were changed to decidualization medium and maintained for 24 hours. MTT solution was added to each well and incubated for 4 hours at 37°C with 5% CO_2_. DMSO was added directly to each well and pipetted up and down to dissolve salts. The absorbance signal was measured on a Perkin Elmer EnVision HTS Multilabel Reader at 570 nm and background absorbance was measured at 630 nm. The optical density was calculated by subtracting the background signal from the absorbance signal.

### RNA extraction and RT-qPCR

Cultured cells were collected and centrifuged at 300 × *g* for 1 minute at 4°C. RNA was isolated using the NucleoSpin RNA extraction kit (Macherey-Nagel, Germany, 740955.50) according to the manufacturer’s protocol. RNA was quantified using an ND-1000 spectrophotometer (NanoDrop). cDNA was prepared from 100 ng of total RNA using iScript cDNA synthesis kit (Bio-Rad, USA, 1708841). Quantitative RT-qPCR was performed in triplicates using SsoAdvanced Universal SYBR Green Supermix, (Bio-Rad, USA, 1725272). The reactions were run in a QuantStudio 6 Flex Real-Time PCR System as follows: 50°C for 2 min and 95°C for 10 min, followed by 40 cycles of 95°C for 15 s and 60°C for 1 min, with a final melt curve step including 95°C for 15 s, 60°C for 1 min, and 95°C for 15 s. Results were normalized to *RPL27*. Primers used in this study are: *CDKN1A*: 5’-AGTCAGTTCCTTGTGGAGCC-3’ and 5’CATTAGCGCATCACAGTCGC--3’; *MRE11*: 5’-CAGCAACCAACAAAGGAAGAGGC-3’ and 5’-GAGTTCCTGCTACGGGTAGAAG-3’; *RAD50*: 5’-GGAAGAGCAGTTGTCCAGTTACG-3’ and 5’-GAGTAAACTGCTGTGGCTCCAG-3’; *NBS1*: 5’-TCTGTCAGGACGGCAGGAAAGA-3’ and 5’-CACCTCCAAAGACAACTGCGGA-3’; *RPL27*: 5’-TGGACAAAACTGTCGTCAATAAGG-3’ and 5’-AGAACCACTTGTTCTTGCCTGTC-3’.

### Single-cell RNA sequencing analysis

The subset stromal tissue anndata object were taken from (Tan et al., 2022) and used in accordance with the authors’ preprocessing and cell type annotations. Analysis focused on healthy control (*n*=3) and endometriosis patient (*n*=9) eutopic samples. Python version 3.12.1 was used with scanpy version 3.12.1 for subsequent analysis (Virshup et al., 2023). Pseudobulk analysis was performed by calculating mean expression of each gene within each patient sample to account for patient-specific effects. Statistical significance was assessed using the Mann-Whitney U test comparing pseudobulk expression values between groups. Effect sizes were calculated using Cohen’s d. For each respective gene analyzed (*NBS1*, *RAD50*, and *MRE11*), cells that had 0 expression were dropped before performing pseudobulk and statistical analysis.

### Western blotting

Cultured cells were collected and centrifuged at 300 × *g* for 1 minute at 4°C. Whole cell lysates were prepared using 1× NP40 lysis buffer (Thermo Scientific, USA, FNN0021) supplemented with Pierce Protease and Phosphatase inhibitors (Thermo Scientific, USA, A32959) and kept on ice for 30 minutes, resuspending every 10 minutes. Lysates were sonicated using a Bioruptor 300 on high for three cycles of 30 sec on and 30 sec off. Protein concentration was quantified using Qubit 4 fluorometer (ThermoFisher, USA, Q33238). Samples were resolved by SDS page and transferred to a polyvinylidene difluoride (PVDF) membrane (Millipore-Sigma, USA, IPVH00010) for 60 minutes at 250 mA. Immunoblots were blocked for 1 hour at room temperature using either 5% non-fat milk or 5% BSA TBS-T. Primary antibodies [anti-β-Actin (SCBT, USA, sc-47778; 1:10,000), anti-p53 (SCBT, USA, sc-126; 1:1000), anti-pATM S1981 (Abcam, United Kingdom, ab81292; 1:1000), anti-ATM (GeneTex, USA, GTX70103; 1:1000), anti-RAD50 (Cell Signaling, USA, 3427T; 1:1000), anti-NBS1 (Cell Signaling, USA, 14956T; 1:1000)] were incubated overnight at 4°C on a rocker. Corresponding HRP-conjugated secondary antibodies (The Jackson Laboratory, USA; 1:5,000) were incubated for 1 hour at room temperature on a rocker. Immunoblots were developed using Clarity ECL (Bio-Rad, USA, 1705060) or Clarity Max ECL (Bio-Rad, USA, 1705062), detected using a Bio-Rad ChemiDoc MP Image, and quantified using ImageJ.

### Clonogenicity assay

Decidualized cells were collected, 1,000 cells were re-plated in a six-well plate in growing medium, and incubated at 37°C with 5% CO_2_. The medium was refreshed every 2-3 days for 12 days. Cell colonies were washed with DPBS, fixed with 4% PFA for 10 minutes, and stained with Crystal violet solution (Sigma, USA, V5265-500ML) for 15 minutes at room temperature. Large, visible colonies were counted.

### Immunocytochemistry (ICC)

Cultured cells in a 96-well plate were fixed at various timepoints with 4% PFA for 10 minutes at room temperature. Cells were washed with 1× PBS and were permeabilized/blocked for 1 hour with 1× PBS containing 10% donkey serum (Equitech-Bio Inc., USA, SD30-0500), 1% BSA (Fisher Bioreagents, USA, BP9703-100), and 0.1% triton (Millipore Sigma, USA, X-100). Cells were washed with 1× PBS and incubated in primary antibodies [anti-vimentin (Biotechne, USA, MAB2105; 1:500), anti-γH2AX S139 (Cell Signaling, USA, 2577S; 1:500), anti-pATM S1981 (GeneTex, USA, GTX132146; 1:500), anti-53BP1 (Cell Signaling, USA, 4937S; 1:500), anti-pCHK2 T68 (Cell Signaling, USA, 2661S; 1:500), anti-p21 (SCBT, USA, sc-6246; 1:500)] in 1× PBS containing 5% donkey serum and 0.1% triton overnight at 4°C. Cells were washed twice with 1× PBS for 5 minutes with gentle shaking. Cells were incubated in secondary antibodies [donkey anti-Rabbit 488 (Invitrogen, USA, XJ357262; 1:500) and donkey anti-Rat 555 (Invitrogen, USA, A21432; 1:500)] and Hoechst 33342 (AnaSpec, USA, AS-83218; 1:1000) in 1× PBS containing 5% donkey serum and 0.1% triton for 2 hours at room temperature in darkness. The cells were washed twice with 1× PBS for 5 minutes with gentle shaking and stored in 1× PBS containing 0.05% sodium azide. Image acquisition was performed on a Perkin Elmer Opera Phenix Plus. Image analysis was performed using CellProfiler. For nuclear integrated intensity, the IdentifyPrimaryObjects module was used to identify individual nuclei and the integrated intensity was measured for each protein. For vimentin integrated intensity, the IdentifyPrimaryObjects module was used to identify individual nuclei, IdentifySecondaryObjects was used to detect vimentin, and the integrated intensity was measured. For foci quantification, individual nuclei and foci were identified using the IdentifyPrimaryObjects module and the RelateObjects module was used to associate the foci with their corresponding nuclei. Focal nuclei were defined as nuclei containing at least one foci and diffuse nuclei had zero foci. The average number of foci per nucleus was calculated by analyzing only the nuclei containing foci while diffuse nuclei were excluded. All integrated intensity measurements were performed on images captured at 20X. All foci and nuclei (eccentricity and area) measurements were performed on images captured at 63X. The number of fields of view and the approximate number of cells or nuclei quantified are denoted in the figure legends.

### SA-β-Gal staining

The senescence-associated β-galactosidase staining kit (Cell Signaling, USA, 9860) was followed according to the manufacturer’s protocol. Cultured cells were fixed using the fixative solution provided for 10 minutes at room temperature. The staining solution was prepared according to the manufacturer’s specifications, and the warmed solution was measured and adjusted as needed to achieve a final pH of 6. The cells were wrapped in parafilm and incubated in the staining solution for 4 hours in a dry 37°C incubator. Cells were washed with 1× PBS and then incubated with Hoechst 33342 (1:1000) for 10 minutes at room temperature. The cells were washed with 1× PBS and then covered in a 70% glycerol (Sigma, USA, G9012) solution. Image acquisition was performed using a Zeiss Live-Cell Imaging Widefield Microscope. The percentage of SA-β-Gal positive cells were quantified using CellProfiler. Individual nuclei were identified using the IdentifyPrimaryObjects module and β-galactosidase positive objects were detected using the IdentifySecondaryObjects. Representative images were acquired using an Echo Revolve microscope.

### Statistical analyses

No statistical method was used to predetermine the sample size. Statistical analysis was performed using Prism10 software or Python for scRNA-seq analysis. Sample size, biological samples, statistical tests, and statistical significance are denoted in the figure legends.

## Results

### Decreased γH2AX Foci in Proliferating MenSCs from Endometriosis Patients

To investigate the DDR, we isolated MenSCs from healthy individuals and those diagnosed with endometriosis (Endo) (Sun, et al., 2019). Cultured MenSCs were collected at three time points to assess the DDR: 1) while proliferating, 2) undecidualized at confluence (UnDec), and 3) after four days of decidualization (Dec) (Figure 1A). We first characterized these cell lines by performing a flow cytometry analysis at the UnDec and Dec timepoints using previously identified markers: CD140b (PDGFRβ), an endometrial stromal cell marker; SUSD2, a marker for perivascular, endometrial stromal cells; and CD146 (MCAM), an endometrial stem/progenitor cell marker (Gargett, et al., 2016, Masuda et al., 2012, Spitzer et al., 2012). There was a high proportion of cells in all the cultures (healthy and Endo, UnDec and Dec) that were CD140b+ and SUSD2+, consistent with their identity as perivascular stromal cells (Suppl Figures 1A-E). We also found that the proportion of CD146+ cells was high in UnDec cultures and reduced after decidualization in both healthy and Endo samples (Figure 1B and Suppl Figure 1F), consistent with these stem/progenitor cells differentiating during decidualization.

**Figure 1:**
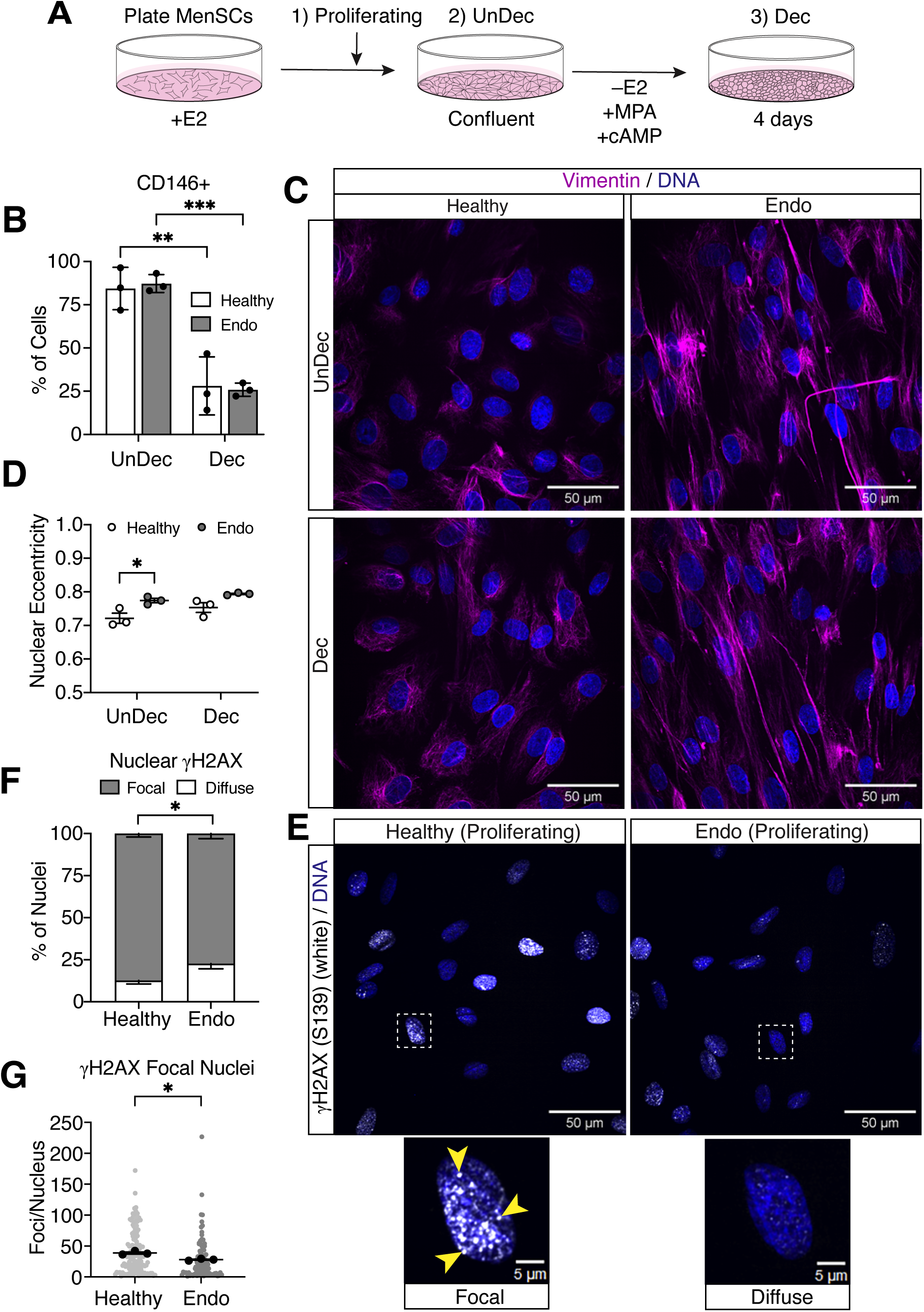
Decreased γH2AX Foci in Proliferating MenSCs from Endometriosis Patients. **A**) Graphical representation of healthy and endometriosis (Endo) menstrual effluent stromal cells (MenSCs) *in vitro* model and collection timepoints: 1) proliferating (80% confluent) in growing medium containing estrogen (E2); 2) 100% confluent, undecidualized (UnDec); and 3) after four days of decidualization (Dec) with cyclic AMP (cAMP) and medroxyprogesterone (MPA). **B**) Percentage of UnDec and Dec, healthy and Endo cells positive for CD146, as detected by flow cytometry. **C**) Representative field of view of UnDec (top) and Dec (bottom), healthy (left) and Endo (right) samples. Vimentin detected in magenta and DNA by Hoechst in blue. **D**) Quantification of UnDec and Dec, healthy and Endo nuclear eccentricity, as detected by ICC. **E**) Representative field of view of γH2AX (white) in proliferating healthy (left) and Endo (right) samples. White dashed lines represent the single magnified nuclei below that indicate focal (left) or diffuse (right) γH2AX. Yellow arrowheads specify foci. DNA by Hoechst in blue. **F**) Percentage of nuclei with focal or diffuse nuclear γH2AX in proliferating healthy and Endo samples. **G**) Quantification of the number of γH2AX foci per focal nucleus in proliferating healthy and Endo samples. For (**D**), dots represent the means of *n*=3 biologically independent samples quantified from 27 fields of view (∼500 nuclei). For (**F**), bars represent the means of *n*=3 biologically independent samples quantified from 9 fields of view (∼100 nuclei). For (**G**), black dots represent the means of *n*=3 biologically independent samples and gray dots represent the individual nuclei of each sample quantified from 9 fields of view (∼100 nuclei). Data representative of *n*=3 biologically independent samples (**B**). Data presented as mean ± SD (**B**, **F**) and mean ± SEM (**D**, **G**). Data analyzed by two-way ANOVA with Tukey’s multiple comparisons test (**B**, **D**) and Welch’s, two-tailed *t* test (**F**, **G**). *p* values: *<0.05, **<0.01, ***<0.001.

To determine if the cultures had morphological differences, we immunostained with anti-vimentin and Hoechst (Figure 1C) and performed shape analyses. Although we saw a variety of vimentin cell shapes, our measurements revealed no significant differences between cultures. In contrast, our analysis of nuclear shape showed more elongated nuclei (higher eccentricity) in UnDec Endo compared to UnDec healthy cells (Figure 1D). Nuclear shape influences cell cycle progression and DDR signaling (Aureille et al., 2019, Dos Santos et al., 2021), suggesting that these functions may be altered in the UnDec Endo samples.

Tissue biopsy studies have consistently reported higher proliferative indices in the eutopic endometrium of women with Endo (Bane, et al., 2021, Hapangama, et al., 2008, Munshi and Sachdeva, 2023, Wingfield et al., 1995). We analyzed cell proliferation by cell cycle analysis via propidium iodide intercalation, Ki67 positivity, and EdU incorporation, and observed no differences (Suppl Figures 1G-K), indicating cell cycle progression and proliferation rates are similar between healthy and Endo MenSCs in culture. These data suggest that proliferative phenotypes in Endo are driven by environmental factors that are not present in our *in vitro* cultures.

We assessed DDR activation during proliferation by immunostaining with anti-γH2AX, a marker of DNA damage, followed by automated high-throughput microscopy. We measured γH2AX intensity (integrated intensity) using a 20X objective, similar to methods used to analyze endometrial tissue biopsies that have shown differences between Endo and healthy tissue (Bane, et al., 2021, Choi, et al., 2018). In contrast, we did not observe differences in γH2AX intensity between healthy and Endo MenSCs (Suppl Figure 1L). We noticed, however, that healthy samples had more nuclei containing punctate γH2AX staining. To investigate, we imaged the samples at high-magnification (63X) and quantified the proportion of nuclei with focal γH2AX staining, while designating nuclei without γH2AX foci as having diffuse staining (Figure 1E). In Endo MenSCs, 77% of the nuclei contained γH2AX foci, whereas in healthy MenSCs, 87% of the nuclei contained foci (Figures 1E, 1F). We then quantified the average number of foci per nucleus and found that Endo nuclei had 27% fewer γH2AX foci compared to healthy nuclei (Figure 1E, 1G). Taken together, these observations indicate that while the DDR is activated in both Endo and healthy MenSCs, the γH2AX signal is propagated less efficiently in proliferating Endo cells, which creates fewer quantifiable foci, possibly due to inadequate recruitment of DDR factors.

### Impaired DNA Damage Response and Attenuated Senescence in MenSCs from Endometriosis Patients

Replication of the genome and faithful repair of DNA damage requires robust activation of the DDR to successfully recruit and retain a multitude of factors responsible for repairing the damage. We assessed DDR activation at confluence in UnDec MenSCs to evaluate unrepaired damage before it is compounded by the chromatin reorganization that occurs during decidualization (Simara et al., 2017, Vrljicak, et al., 2018). As we observed in proliferating MenSCs, the intensity of γH2AX was similar between UnDec healthy and Endo samples, but there was no difference in the proportion of nuclei with focal γH2AX staining (Suppl Figures 2A, 2B). However, we found that Endo samples had 19% fewer foci per nucleus than healthy samples (Figures 2A, 2B). This recapitulates the reduced γH2AX foci number observed in Endo samples during proliferation, again suggesting a defective DDR that is unable to adequately propagate the γH2AX signal.

**Figure 2:**
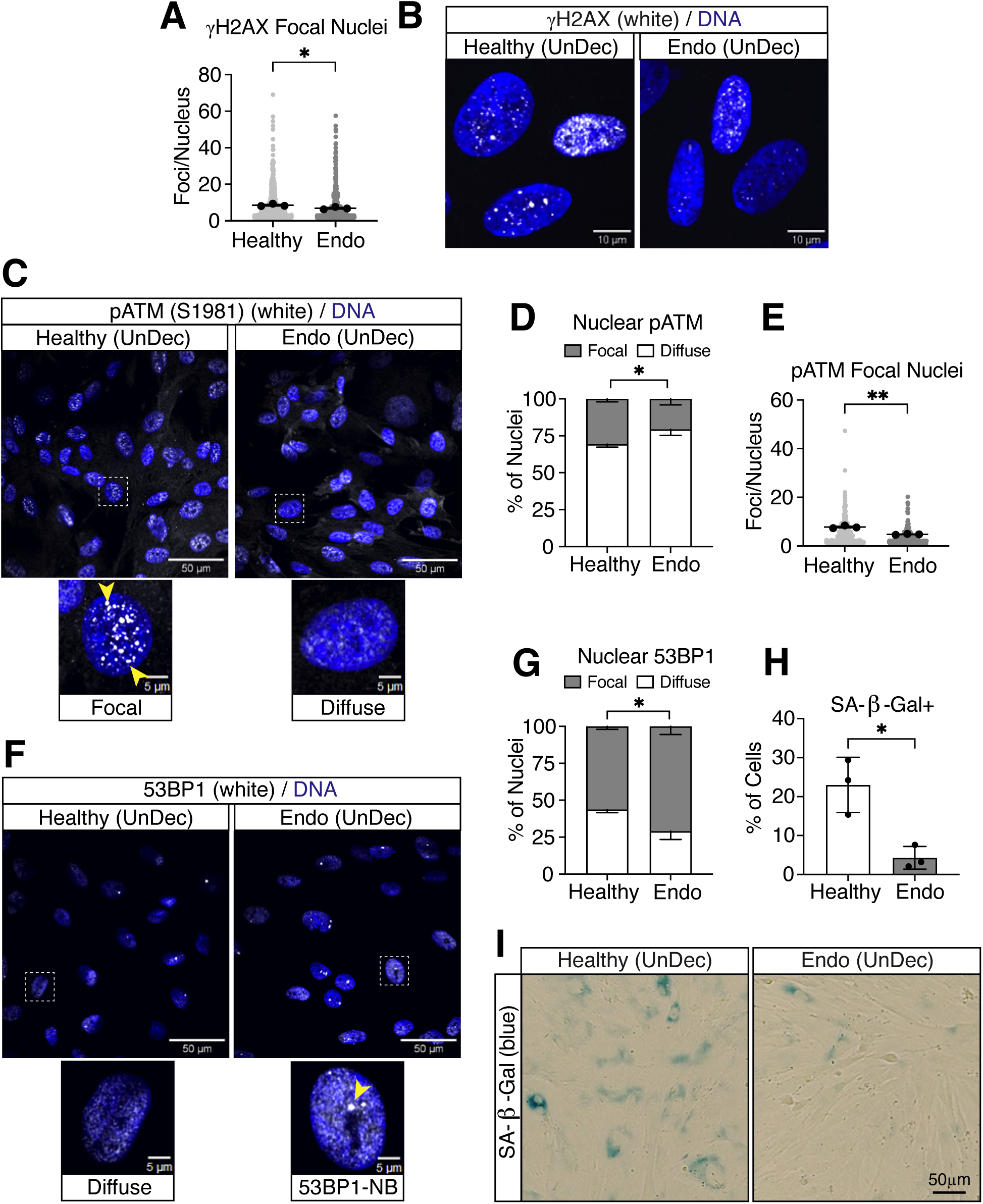
Impaired DNA Damage Response and Attenuated Senescence in MenSCs from Endometriosis Patients. **A**) Quantification of the number of γH2AX foci per focal nucleus in UnDec healthy and Endo samples. **B**) Representative images of γH2AX (white) in UnDec healthy (left) and Endo (right) nuclei. DNA by Hoechst in blue. **C**) Representative field of view of pATM (white) in UnDec healthy (left) and Endo (right) nuclei. White dashed box represents the single magnified nuclei shown below that indicate focal (left) or diffuse (right) pATM. Yellow arrowheads specify foci. DNA by Hoechst in blue. **D**) Percentage of nuclei with focal or diffuse nuclear pATM in UnDec healthy and Endo samples. **E**) Quantification of the number of pATM foci per focal nucleus in UnDec healthy and Endo samples. **F**) Representative field of view of 53BP1 (white) in UnDec healthy (left) and Endo (right) nuclei. White dashed box represents the single magnified nuclei shown below that indicate diffuse (left) or focal 53BP1-NB (right) 53BP1. Yellow arrowhead specifies 53BP1-NB foci. DNA by Hoechst in blue. **G**) Percentage of nuclei with focal or diffuse nuclear 53BP1 in UnDec healthy and Endo samples. **H**) Percentage of cells that are positive for SA-β-Gal staining in UnDec healthy and Endo samples. **I**) Representative images of SA-β-Gal staining (blue) from UnDec healthy (left) and Endo (right) samples. For (**A**, **E**), black dots represent the means of *n*=3 biologically independent samples and gray dots represent the individual nuclei of each sample quantified from 27 fields of view (∼1,000 nuclei). For (**D**, **G**), bars represent the means of *n*=3 biologically independent samples quantified from 27 fields of view (∼1,000 nuclei). Data representative of *n*=3 biologically independent samples (**H**). Data presented as mean ± SD (**D**, **G**, **H**) and mean ± SEM (**A, E**). Data analyzed by Welch’s, two-tailed *t* test (**A**, **D**, **E**, **G**, **H**). *p* values: *<0.05, **<0.01.

γH2AX foci have been used as a DSB biomarker (Redon et al., 2011), therefore we evaluated pATM, which is responsible for phosphorylating γH2AX at DSB sites. To assess ATM activation, we quantified pATM intensity and found it was similar between healthy and Endo samples (Suppl Figure 2C). pATM also localizes into visible foci when it is robustly recruited and sufficiently retained at the DSB, allowing for successful repair of the damage (So et al., 2009). We quantified the proportion of nuclei with focal ATM staining, again designating nuclei without pATM foci as diffuse (Figure 2C). There were 10% fewer Endo nuclei with pATM foci (Figures 2C, 2D). We also observed 37% fewer foci per nucleus in Endo samples (Figure 2C, 2E), which is consistent with the reduced γH2AX foci number in these samples (Figures 1G, 2A). This reduction in recruitment and retention of pATM into foci in Endo samples may result in unrepaired DSBs and/or prolonged repair of DBSs.

Cells with large genomes, such as human MenSCs, will fail to replicate all of their DNA during S-phase, resulting in regions of under-replicated DNA that persist beyond mitosis (Al Mamun et al., 2016, Moreno et al., 2016). Some of these regions contain DSBs that are protected and marked by 53BP1-NBs in G1-phase of the cell cycle (Harrigan, et al., 2011, Lukas, et al., 2011). To determine the cell cycle phase of confluent UnDec MenSCs, we performed cell cycle analysis and found that the majority of cells (86%) were in G1/G0 (Suppl Figure 2D), suggesting these cells contain 53BP1-NBs. We quantified the proportion of nuclei with 53BP1-NB foci, again designating nuclei without foci as diffuse (Figure 2F). Endo samples had 13% more nuclei containing 53BP1-NBs than healthy samples (Figure 2F, 2G), although the number of NBs per nucleus was similar between these samples (Suppl Figure 2F). We also measured 53BP1 intensity by immunostaining in UnDec MenSCs and found similar levels in healthy and Endo samples (Suppl Figure 2E), reminiscent of the intensity results for γH2AX and pATM (Suppl Figures 2A, 2C). Altogether, these data suggest that the localization of DDR factors, γH2AX and pATM, into foci is responsible for driving the DDR defect in Endo MenSCs, resulting in more cells containing under-replicated DNA regions and unrepaired DSBs in 53BP1-NBs.

Previous studies have shown that accumulation of unrepaired DSBs in γH2AX foci results in senescence (Sedelnikova et al., 2004). To determine if this occurred in our cultures, we performed senescence-associated β-galactosidase (SA-β-Gal) assays. In healthy samples, 23% of cells stained positive for SA-β-Gal, consistent with a recent study examining primary endometrial stromal cells from tissue biopsies (Deryabin and Borodkina, 2022), compared to only 4% of cells in the Endo samples (Figures 2H, 2I). This suggests that Endo cells are less prone to senescence when proliferating to confluence; however, it seemed unlikely that the Endo samples accrued significantly less DNA damage than healthy samples. Instead, our data suggest that a defective DDR in Endo samples does not allow for the proper recognition of DSBs and the robust signaling through ATM that is required to induce senescence (Sedelnikova, et al., 2004).

### Replication Stress Increases Unrepaired DNA Damage in MenSCs from Endometriosis Patients

A highly proliferative tissue, such as the endometrium, is subject to replication stress, which is defined as barriers that interfere with DNA replication and can lead to DNA damage (Gaillard et al., 2015). To explore the response of cultured MenSCs to replication stress, we treated proliferating cells with hydroxyurea (Hu), which inhibits nucleoside synthesis. We performed a Hu titration analysis and MTT assays to ensure that none of the selected Hu dosages impaired the viability or metabolic activity of the cells (Suppl Figures 3A, 3B). DDR activation was then assessed by immunostaining with anti-γH2AX when the cells reached confluence. As expected in UnDec samples, increasing doses of Hu resulted in increased γH2AX intensity in both healthy and Endo samples compared to untreated samples. At higher Hu concentrations, however, Endo nuclei consistently exhibited greater γH2AX intensity than healthy nuclei (Figure 3A). Using the highest concentration of Hu treatment (10mM) that did not affect cell viability, we observed a significant increase in γH2AX foci size in Endo nuclei (Figures 3B, 3D), as well as a trending increase in the number of γH2AX foci per nucleus (Figures 3C, 3D). There was no difference in the portion of nuclei containing γH2AX foci between healthy and Endo treated samples (Suppl Figure 3C). This suggests that Endo samples have increased activation of the DDR and more damage marked by γH2AX after experiencing replication stress from a high dosage of Hu treatment. These results contrast with our data on the untreated samples where Endo samples had fewer γH2AX foci (Figures 1G, 2A); however, they are more similar to the observations on Endo tissue, where higher levels of γH2AX were observed (Bane, et al., 2021, Choi, et al., 2018).

**Figure 3:**
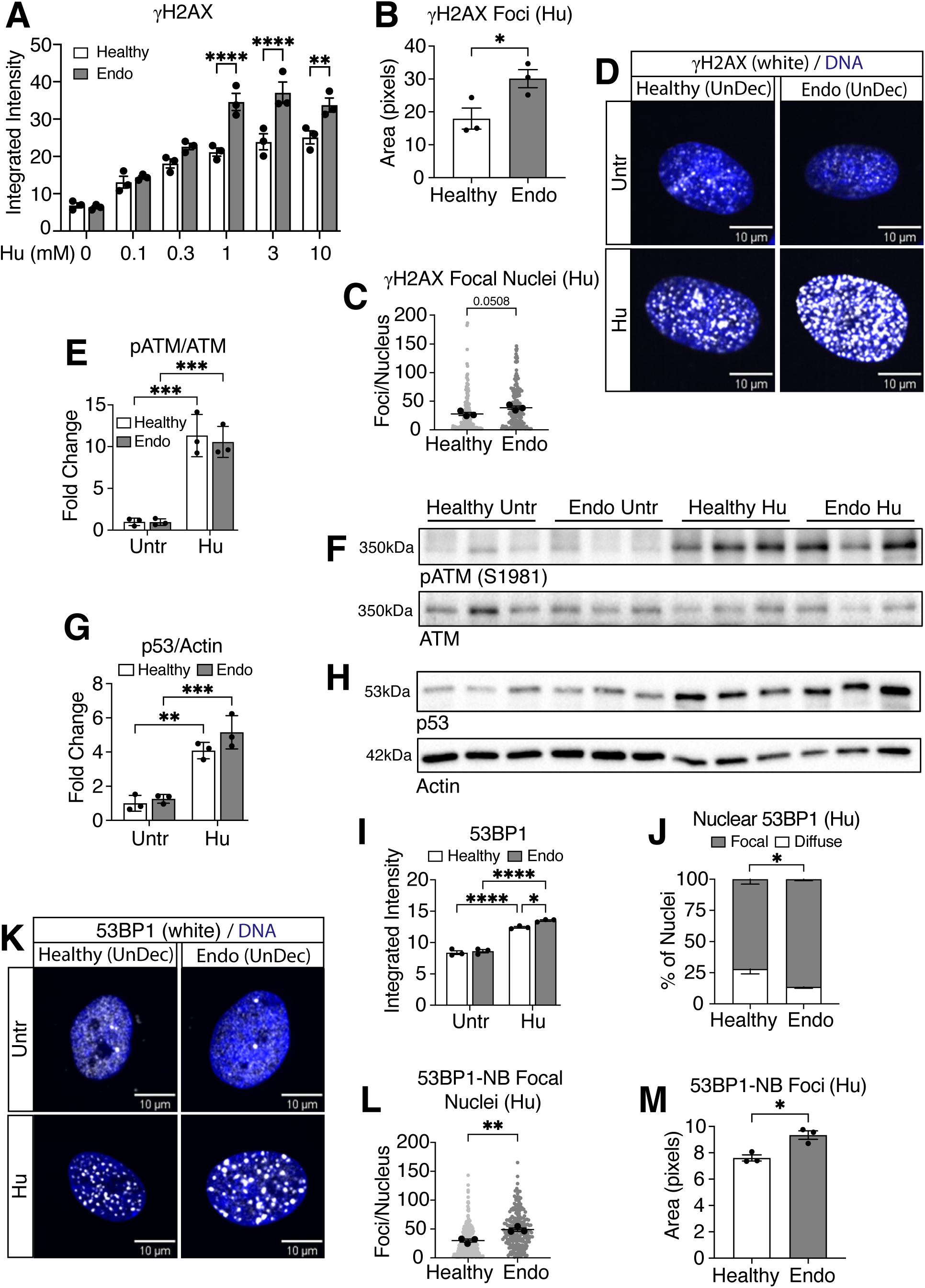
Replication Stress Increases Unrepaired DNA Damage in MenSCs from Endometriosis Patients. **A**) Quantification of nuclear γH2AX integrated intensity in untreated and Hu-treated, UnDec healthy and Endo samples, as detected by ICC. **B**) Quantification of γH2AX foci area in Hu-treated, UnDec healthy and Endo samples. **C**) Quantification of the number of γH2AX foci per focal nucleus in Hu-treated, UnDec healthy and Endo samples. **D**) Representative images of γH2AX (white) from untreated (top) and Hu-treated (bottom), UnDec healthy (left) and Endo (right) samples. DNA by Hoechst in blue. **E**, **F**) Quantification (**E**) and western blot (**F**) of pATM expression in untreated and Hu-treated, UnDec healthy and Endo samples. Quantification shown as pATM/ATM ratio. **G**, **H**) Quantification (**G**) and western blot (**H**) of p53 expression in untreated and Hu-treated, UnDec healthy and Endo samples. Normalized to Actin loading control. **I**) Quantification of nuclear 53BP1 integrated intensity in untreated and Hu-treated, UnDec healthy and Endo samples, as detected by ICC. **J**) Percentage of nuclei with focal or diffuse nuclear 53BP1 in Hu-treated, UnDec healthy and Endo samples. **K**) Representative images of 53BP1 (white) from untreated (top) and Hu-treated (bottom), UnDec healthy (left) and Endo (right) samples. DNA by Hoechst in blue. **L**) Quantification of the number of 53BP1-NB foci per focal nucleus in Hu-treated, UnDec healthy and Endo samples. **M**) Quantification of 53BP1-NB foci area in Hu-treated, UnDec healthy and Endo samples. For (**C**, **L**), black dots represent the means of *n*=3 biologically independent samples and gray dots represent the individual nuclei of each sample quantified from 27 fields of view (∼150 nuclei). For (**J**), bars represent the means of *n*=3 biologically independent samples quantified from 27 fields of view (∼150 nuclei). Data representative of *n*=3 biologically independent samples (**A**, **B**, **E**, **F**, **G**, **H**, **I**, **M**). Data presented as mean ± SD (**E**, **G**, **J**) and mean ± SEM (**A**, **B**, **C**, **I**, **L**, **M**). Data analyzed by two-way ANOVA with Šídák’s (**A**) or Tukey’s (**E**, **G**, **I**) multiple comparison test and Welch’s, two-tailed *t* test (**B, C, J, L, M**). *p* values: *<0.05, **<0.01, ***<0.001, ****<0.0001.

To evaluate the mechanism driving increased γH2AX intensity in Hu-treated Endo samples, we examined the ATM-dependent DDR pathway. As expected, we found upregulation of pATM in response to Hu treatment in both healthy and Endo samples (Figures 3E, 3F and Suppl Figure 3D, 3E). Remarkably, we did not observe a concomitant increase in pATM in treated Endo nuclei compared to healthy nuclei, as measured by the proportion of nuclei with foci, the number of foci per nucleus, or the foci area (Suppl Figure 3E-H), despite the increased γH2AX intensity in Endo samples (Figure 3A). We also examined the expression of downstream effectors, pChk2, p53, and p21, and observed similar increases in expression of each one in response to Hu treatment, but again there were no differences between healthy and Endo Hu-treated samples (Figures 3G, 3H and Suppl Figures 3I-M). These data suggest that replication stress promotes checkpoint activation in cultured MenSCs, allowing both Endo and healthy cells to recognize and mark damaged DNA with γH2AX: a process that did not efficiently occur in Endo samples in the absence of replication stress (Figures 1G, 2A). In addition, ATM-dependent pathway activation was not greater in Hu-treated Endo samples than treated healthy samples, suggesting that the defective DDR in Endo samples leads to persistent γH2AX signals but inadequate pathway activation needed to repair the damage.

To investigate the unrepaired damage from replication stress in MenSCs, we analyzed 53BP1 by immunostaining and observed increased intensity with Hu treatment in both healthy and Endo samples (Figure 3I), consistent with increased replication stress. In Hu-treated samples, however, there is greater 53BP1 intensity in Endo nuclei compared to healthy nuclei (Figure 3I). We also found that treated Endo samples had 14% more nuclei with 53BP1-NBs, 64% more 53BP1-NBs per nucleus, and 22% larger NBs (Figures 3J-M). These data suggest that replication stress in Endo samples results in more unrepaired damage and regions of under-replicated DNA due to insufficient activation of repair pathways; nevertheless, the cells contain the damage in 53BP1-NBs in order to counteract their defective DDR.

### MenSCs from Endometriosis Patients Exhibit Reduced Expression of the MRN Complex

The initial detection of DSBs is one of the crucial roles of the MRN complex, which detects the broken ends of DSBs and activates the DDR by facilitating ATM autophosphorylation. It has been shown that MRN deficiency in S-phase increases the burden of replication-associated lesions carried into G1, manifesting as elevated numbers of 53BP1-positive nuclear bodies (Kondratova et al., 2015); although, compositions of these nuclear bodies may be altered based on the availability of repair factors (Fernandez-Vidal et al., 2017). Thus, one explanation for the increase in 53BP1-NBs observed in Endo samples (Figures 3J-M) is defects in the MRN complex. To investigate, we used RT-qPCR to examine the expression of the MRN complex components, *MRE11*, *RAD50*, and *NBS1*, in our samples. While no significant differences were observed in untreated samples, Hu treatment resulted in significant reductions in expression of each gene in Endo samples compared to healthy samples (Figure 4A).

**Figure 4:**
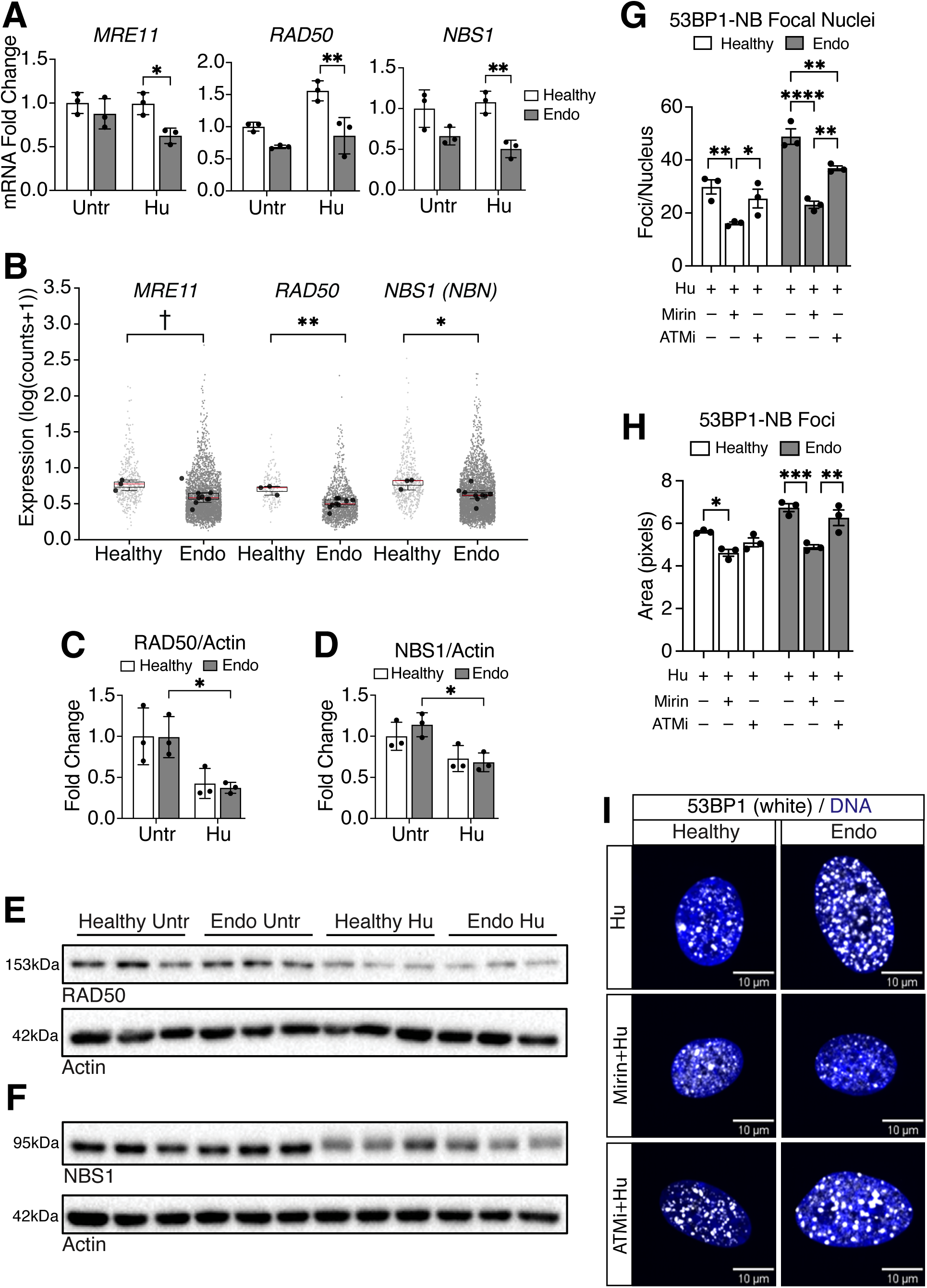
MenSCs from Endometriosis Patients Exhibit Reduced Expression of the MRN Complex. **A)** MRN complex gene expression, *MRE11*, *RAD50*, and *NBS1*, in untreated and Hu-treated, UnDec healthy and Endo samples, as detected by RT-qPCR. **B**) MRN complex gene expression, *MRE11*, *RAD50*, and *NBS1* (*NBN*), from a single cell RNA-seq data set of healthy control and Endo samples. Gray dots represent expression of each cell analyzed from each sample and the black dots represent the pseudobulk expression for *n*=3 healthy control samples and *n*=9 Endo patient samples. Data presented as a boxplot of the pseudobulk expression and the red bar represents the median expression. **C-F**) Quantification and western blot of RAD50 (**C**, **E**) and NBS1 (**D**, **F**) expression in untreated and Hu-treated, UnDec healthy and Endo samples. Normalized to Actin loading control. **G**) Quantification of the number of 53BP1-NB foci per focal nucleus in Hu-, Mirin+Hu-, and ATMi+Hu-treated, UnDec healthy and Endo samples. **H**) Quantification of 53BP1-NB foci area in Hu-, Mirin+Hu-, and ATMi+Hu-treated, UnDec healthy and Endo samples. **I**) Representative images of 53BP1 (white) from Hu-treated (top), Mirin+Hu-treated (middle), and ATMi+Hu-treated (bottom), UnDec healthy (left) and Endo (right) samples. DNA by Hoechst in blue. Data representative of *n*=3 biologically independent samples (**A**, **C**-**F**). For (**G**, **H**), data represents the means of *n*=3 biologically independent samples quantified from 27 fields of view (∼150 nuclei). Data presented as mean ± SD (**A**, **C**, **D**) and mean ± SEM (**G**, **H**). Data analyzed by two-way ANOVA with Tukey’s multiple comparison test (**A**, **C**, **D**, **G**, **H**) and the Mann-Whitney U test comparing pseudobulk expression values between groups (**B**). *p* values: †=0.06, *<0.05, **<0.01, ***<0.001, ****<0.0001.

To validate the RT-qPCR findings *in vivo*, we interrogated a scRNA-seq dataset for gene expression of *MRE11*, *RAD50*, and *NBS1* in the stromal subpopulation of eutopic endometrial tissue from healthy and Endo samples (Tan, et al., 2022). We found a trending decrease in *MRE11* and significant reductions in *RAD50* and *NBS1* in endometriosis patients (Figure 4B). We also examined MRN complex proteins in our cultured MenSCs by immunoblot and found reduced protein expression of RAD50 and NBS1 in Hu-treated Endo samples, whereas the reductions of these proteins in healthy treated samples were not significant (Figures 4C-F). These data again indicate that Endo samples are under replication stress *in vivo* and suggests that their defective response to DNA damage and replication stress is due to a MRN deficiency, leading to insufficient detection of DSBs in S-phase, inadequate activation of the DDR, and incomplete repair.

We observed that the expression of each component of the MRN complex is reduced in Hu-treated Endo samples. To evaluate reduced MRN pathway activity in healthy cells, we used two drugs: Mirin, which blocks MRN’s ability to activate ATM (Dupre et al., 2008); and the ATM inhibitor, Ku60019 (ATMi) that blocks ATM kinase activity and auto-phosphorylation at Ser1981 (Golding et al., 2009). We evaluated the effect of MRN or ATM inhibition under the condition of replication stress, using 53BP1 foci formation as a readout. As previously observed, Hu treatment, alone, resulted in more and larger 53BP1-NBs in Endo samples (Figure 3L, 3M). Treating with both Hu and Mirin resulted in a reduction of 53BP1-NB foci number and size in both healthy and Endo samples compared to Hu treatment alone (Figures 4G-I). Treating with both Hu and ATMi, however, resulted in reduced 53BP1-NB foci number only in Endo samples compared to Hu treatment alone, with foci size remaining similar (Figures 4G-I). These data suggest that Endo MenSCs are particularly sensitive to pharmacological inhibition of ATM (Figures 4G-I), likely due to their concurrent reduced MRN expression (Figures 4A, 4C-F).

Furthermore, interpreting the results from this pharmacological inhibitor experiment led to an interesting finding because we might have expected to see more 53BP1-NBs in both healthy and Endo samples after inhibiting the MRN complex, due to cells accumulating replication-associated lesions in the absence of MRN activity. Instead, we observed a decrease in the number of 53BP1-NBs in both samples because there is a difference between reduced expression (Figures 4A, 4C-F) and pharmacological inhibition (Figures 4G-I). Pharmacological inhibition of MRN throughout the entire cell cycle reduces 53BP1-NB formation during G1 in both Endo and healthy samples (Figures 4G-I), because both MRN and ATM are components of 53BP1-NBs (Fernandez-Vidal, et al., 2017). Thus, pharmacological inhibition leads to a different outcome in cells compared to reduced MRN complex expression, with these inhibitor studies supporting a model whereby reduced MRN complex expression is only detrimental during S-phase, when many MRN complexes are needed to initiate DNA repair. MRN reduction in this context results in insufficient DDR activation and unrepaired DSBs. In the absence of inhibitors, Endo MenSCs are able to tolerate this reduction by forming “catch-up” 53BP1-NBs in G1 (Figures 3J-M), when Endo cells have more time to deploy their diminished MRN to damage sites and activate ATM. Taken together, these data suggest that in Endo samples MRN insufficiently activates the DDR during S-phase to repair damage; however, MRN-ATM activity is sufficient during G1 to form 53BP1-NBs.

### Subpopulation of MenSCs from Endometriosis Patients Fail to Undergo Senescence upon Decidualization and Instead Retain Clonogenicity

During the menstrual cycle, stromal cells undergo decidualization post-ovulation in a process that can be recapitulated *in vitro* (Doi-Tanaka et al., 2024, Sugawara et al., 2014). To assess unrepaired DNA damage from replication stress and from decidualization in our MenSCs cultures, we analyzed the number of γH2AX foci and 53BP1-NBs per nucleus in untreated and Hu-treated Dec samples. While we found no difference in the number of γH2AX foci per nucleus between untreated healthy and Endo samples, there was a greater number in both samples after Hu-treatment (Figures 5A, 5B). We further observed that Hu-treated Endo nuclei contained 47% more γH2AX foci per nucleus than treated healthy nuclei (Figures 5A, 5B). This is similar to the results seen in UnDec Hu-treated Endo samples (Figures 3A-D). Next, we quantified 53BP1-NBs and observed similar numbers of 53BP1-NBs per nucleus in untreated healthy and Endo cells (Figures 5C, 5D). In contrast, Hu-treated Endo samples have more NBs per nucleus and more cells with 53BP1-NBs following decidualization than both their untreated control samples and Hu-treated healthy samples (Figures 5C-E). Together, these data suggest that replication stress and decidualization lead to more unrepaired and persistent DNA damage in Endo samples.

**Figure 5:**
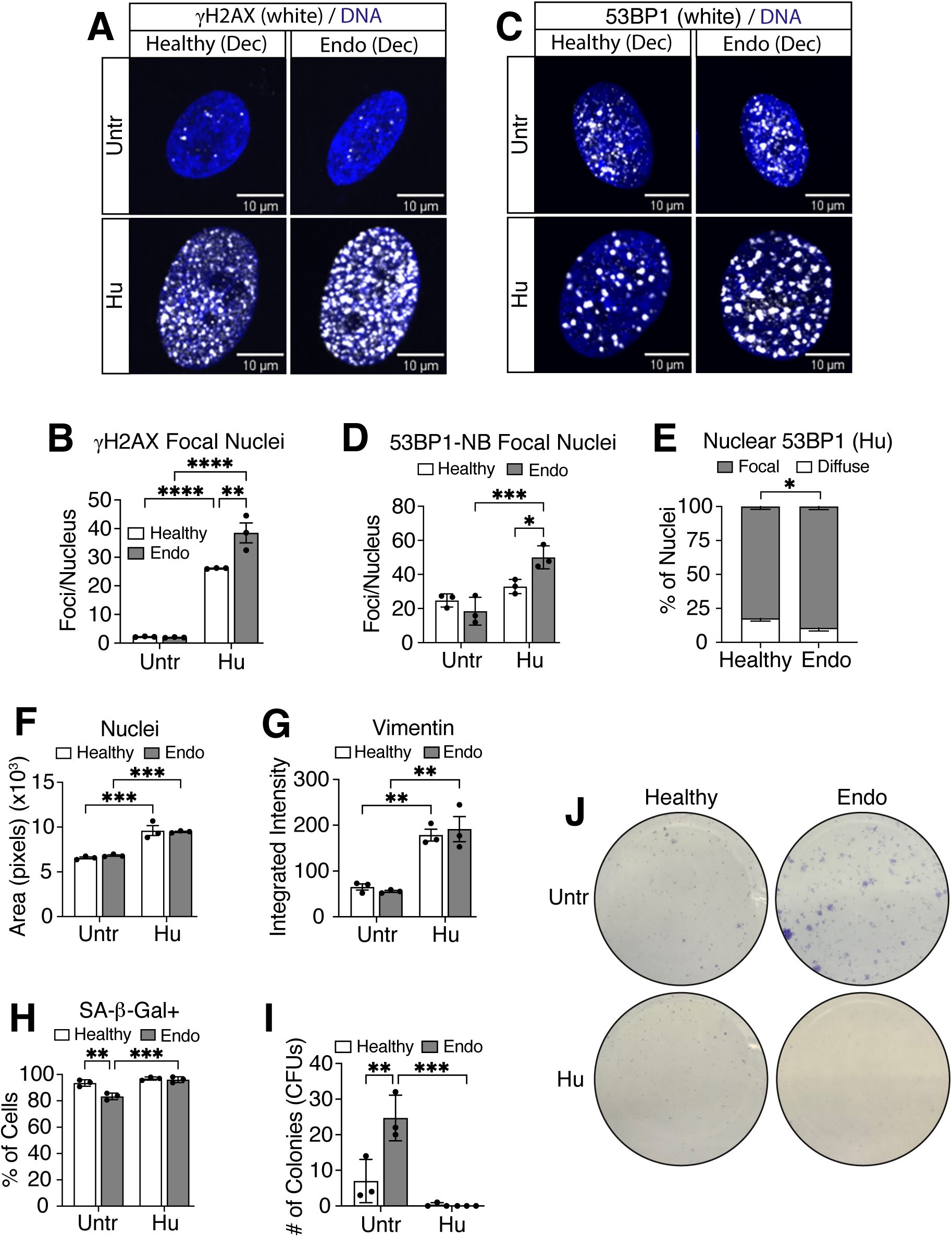
Subpopulation of MenSCs from Endometriosis Patients Fail to Undergo Senescence upon Decidualization and Instead Retain Clonogenicity. **A)** Representative images of ãH2AX (white) from untreated (top) and Hu-treated (bottom), Dec healthy (left) and Endo (right) nuclei. DNA by Hoechst in blue. **B**) Quantification of the number of γH2AX foci per focal nucleus in untreated and Hu-treated, Dec healthy and Endo samples. **C**) Representative images of 53BP1 (white) from untreated (top) and Hu-treated (bottom), Dec healthy (left) and Endo (right) samples. DNA by Hoechst in blue. **D**) Quantification of the number of 53BP1-NB foci per focal nucleus in untreated and Hu-treated, Dec healthy and Endo samples. **E**) Percentage of nuclei with focal or diffuse nuclear 53BP1 in Hu-treated, Dec healthy and Endo samples. **F**) Quantification of nuclei area in untreated and Hu-treated, Dec healthy and Endo samples, as detected by ICC. **G**) Quantification of vimentin integrated intensity in untreated and Hu-treated, Dec healthy and Endo samples, as detected by ICC. **H**) Percentage of cells positive for SA-β-Gal staining in untreated and Hu-treated, Dec healthy and Endo samples. **I)** Quantification of colony forming units from clonogenicity assay of untreated and Hu-treated, Dec healthy and Endo samples. **J**) Representative images of clonogenicity assay from untreated (top) and Hu-treated (bottom), Dec healthy (left) and Endo (right) samples. For (**B**, **D**, **E**, **F**) data represents the means of *n*=3 biologically independent samples quantified from 27 fields of view (untreated ∼500 nuclei; Hu ∼150 nuclei), or for (**G**), 9 fields of view (∼1,000 cells). Data representative of *n*=3 biologically independent samples (**H**, **I**). Data presented as mean ± SD (**E**, **H**, **I**) and mean ± SEM (**B**, **D**, **F**, **G**). Data analyzed by two-way ANOVA with Tukey’s multiple comparison test (**B**, **D**, **F**, **G**, **H**, **I**) and Welch’s, two-tailed *t* test (**E**). *p* values: *<0.05, **<0.01, ***<0.001, ****<0.0001.

Previous studies have shown that decidualization also induces a senescent subpopulation of stromal cells in primary cultures of endometrial tissue biopsies (Brighton, et al., 2017). To determine the effects of decidualization and replication stress on senescence in MenSCs, we first measured nuclei area and vimentin intensity, markers of senescence, and observed that Hu treatment increased both of these markers in decidualized healthy and Endo samples (Figures 5F, 5G and Suppl Figure 4A). We then evaluated SA-β-Gal activity and found that the proportion of Dec MenSCs staining positive was high in healthy samples, regardless of treatment (Figure 5H and Suppl Figure 4B). In contrast, the proportion of positive cells was 10% lower in untreated Dec Endo samples, similar to our results in UnDec Endo samples (Figures 2H, 2I). Nevertheless, Hu treatment increased the proportion of senescent Dec Endo cells up to the levels seen in Dec healthy cells (Figure 5H and Suppl Figure 4B). This again indicates that Endo samples are less prone to senescence, even though replication stress and DNA damage promote senescence.

Shed endometrial cells are exposed to the body cavity through retrograde menstruation, where they might attach and proliferate, creating ectopic lesions. To explore the proliferative capacities of the Dec MenSCs, we performed clonogenicity assays by trypsinizing the cells and replating them in growing medium. We found that untreated Dec healthy MenSCs grew few colonies; in contrast, untreated Dec Endo MenSCs grew 3.5-fold more colonies (Figure 5I, 5J). This suggests the untreated Dec Endo MenSCs retained their proliferative capacity, which is consistent with our observations that fewer untreated UnDec and Dec Endo cells senesced (Figure 2H, 5H). Hu-treated cells from both healthy and Endo cultures had little to no proliferative capacity (Figure 5I, 5J). Taken together, our results suggest that in some patients with endometriosis, there is an inability of eutopic MenSCs to mount an effective DDR, preventing senescence and leaving a small percentage of cells with proliferative capacity that could contribute to the development of ectopic endometrial lesions. Replication stress exacerbates their defective DDR, impeding complete repair of damaged DNA and promoting senescence.

## Discussion

One fundamental question in the field of endometriosis is: How do ectopic endometrial cells persist as lesions? They are not cancerous, per se, yet they are able to proliferate and invade, instead of apoptosing or fully senescing, as would be expected of non-cancerous cells that find themselves in a foreign niche. Generally, cells face environmental and architectural control over their proliferation, but in endometriosis patients, endometrial stromal and epithelial cells in alternative microenvironments continue to thrive. These findings point to inherent cellular characteristics within the endometriosis-derived endometrium that enable survival, implantation, and expansion at ectopic sites. At the same time, the widely recognized anatomical, histological, and molecular heterogeneity of endometriosis suggests that multiple mechanisms contribute to its pathogenesis, which in turn, complicates efforts to fully define its etiology and to develop comprehensive diagnostic and therapeutic strategies.

Retrograde menstruation, in which viable endometrial tissue refluxes through the fallopian tubes into the peritoneal cavity during menses, remains a widely favored theory for the pathogenesis of endometriosis (Zondervan, et al., 2020). In line with this hypothesis, we examined the DNA damage response in menstrual effluent-derived stromal cells (MenSCs), which serve as a model for investigating the properties of endometrial cells that may implant ectopically and drive disease. Our results corroborate earlier studies on the DDR in eutopic endometrial tissue biopsies (Bane, et al., 2021, Choi, et al., 2018). Bane et al. discovered a higher immunostaining score of nuclear γH2AX foci-positive cells in the stromal compartment in Endo compared to healthy samples (Bane, et al., 2021), with a similar finding by Choi et al. (Choi, et al., 2018), suggesting an upregulation of the DDR in response to replication and other stresses, such as oxidative and inflammatory, that are observed in tissue samples from endometriosis patients (Munshi and Sachdeva, 2023). We also observed a significant increase in γH2AX intensity as well as foci size in Endo samples, but only those subjected to replication stress. Thus, in our *in vitro* MenSC cultures, Hu-treated Endo samples more closely mirror the findings of increased DDR activation in tissue biopsies, reinforcing the notion that patient tissue has experienced substantial stress and defective DNA damage repair (Munshi and Sachdeva, 2023).

In search of a molecular explanation for the defective DDR observed in our Endo MenSCs, we identified reduced expression of genes encoding the MRN complex, *MRE11*, *RAD50* and *NBS1*, in our three patient-derived cell lines and in a scRNA-seq dataset of eutopic endometrium biopsies from nine Endo patients. MRN sits at the top of the DSB response: it recognizes the broken DNA, holds the two DNA ends together, and switches on ATM to launch checkpoint signaling and repair. Given such a key role in DSB repair, it is not surprising that inherited mutations in these MRN genes would have serious consequences (McCarthy-Leo et al., 2022). Germline mutations in *MRE11*, *RAD50*, and *NBS1* cause Ataxia-telangiectasia-like disease (ATLD), Nijmegen breakage syndrome (NBS), and NBS-like disorder (NBSLD). There is also the related ataxia-telangiectasia (A-T) syndrome that is caused by mutations in the ATM kinase. Mutations or loss-of-function in MRN complex genes are also found in a variety of human cancers and have been linked to increased sensitivity to DNA damage-targeted therapies. For example, MRE11-deficient endometrial cancers showed heightened PARP-inhibitor sensitivity (Koppensteiner et al., 2014), *RAD50*-mutant tumors with compromised ATM signaling demonstrated synthetic lethality when treated with a combination of CHK1 inhibitors and DNA-damaging chemotherapy (Al-Ahmadie et al., 2014), and germline *NBS1* mutation in a uterine sarcoma rendered a sustained response to PARP inhibitory therapy (Markert et al., 2022). In these genetic and disease contexts, the loss or reduction of one MRN complex member frequently results in the reduced steady-state levels of the others, consistent with a requirement for inter-subunit interactions for MRN stability (Kim et al., 2017, von Morgen et al., 2017).

Catastrophic loss of MRN activity, however, does not model what we observed. In our three patient-derived cell lines we found modest reductions in expression of MRN complex genes and proteins. We found this led to defective DSB DDR activation in S-phase, resulting in the formation of “catch-up” 53BP1-NBs in G1, marking regions of under-replicated and unrepaired DNA damage. We also observed reduced senescence in the Endo samples that had not been subjected to replication stress. This observation is in line with MRN deficiency preventing efficient ATM activation and ATM-dependent checkpoint signaling (Kondratova, et al., 2015). Defective MRN-ATM signaling abrogates checkpoint enforcement and the senescence response to DSBs (Di Micco et al., 2006). This defect in MenSC senescence has important implications for endometriosis disease progression as previous studies have also shown decreased decidualization of endometrial stomal cells (Barragan et al., 2016). The deficit in decidualization combined with reduced senescence sets the stage for the growth of shed MenSCs in the pelvic cavity. Indeed, we found that upon decidualization and without replication stress, Endo samples were also more clonogenic, evidence that Endo MenSCs, with their diminished MRN-ATM signaling, may have a population of cells with proliferative potential. Replication stress, however, amplified the γH2AX signal in these Endo cell lines despite diminished MRN, likely due to the activation of other DNA repair kinases, and prolonged or incomplete repair of damage (Furuta et al., 2003, Stiff et al., 2004, Tomimatsu et al., 2009). Hu-treated Endo samples did not retain clonogenicity upon decidualization, suggesting replication stress could heighten the sensitivity of these cells to their MRN deficiency, as observed in some cancers, and prevent their ability to grow outside the uterus (Al-Ahmadie, et al., 2014, Koppensteiner, et al., 2014, Markert, et al., 2022). In sum, our data supports a model in which MRN-ATM deficiency in endometriosis MenSCs produces both proliferative and highly damaged cells after decidualization.

There are limitations inherent in our study. First, we used MenSCs to model cells that may escape into the peritoneal cavity, but this is an *in vitro* model with MenSC lines generated, frozen, and subjected to limited passage. MenSCs were obtained from healthy donors and endometriosis patients who reported their status. While we requested samples from endometriosis patients who were not taking birth control, they had varying disease stages and medical histories. Moreover, the non-endometriosis status of the control samples was not confirmed surgically. A second limitation of our study is the small sample size of three healthy and three Endo MenSC lines. This is mitigated by scRNA-seq analysis that confirmed our finding that MRN complex genes are reduced in eutopic endometriosis samples of nine additional patients, but nonetheless our findings should be verified on larger cohorts. Future investigations assessing markers such as 53BP1 directly in endometrial tissue specimens, both eutopic and ectopic, would constitute an important step toward validating and extending our *in vitro* observations. In addition, our data demonstrate that transcripts encoding several MRN complex components are downregulated in individuals with endometriosis, underscoring the need to elucidate the regulatory mechanisms responsible for this alteration. One plausible mechanism is DNA-damage-induced transcriptional stress, which has been proposed to cause global reductions in essential mRNAs (Lans et al., 2019).

In brief, our observations show that proliferating Endo MenSCs display a defective DDR. This does not necessarily mean that endometriosis cells incur less DNA damage; rather, our data suggest their defective DDR compromises proper DSB recognition and repair activation, leading to more under-replicated DNA, less senescence, and proliferative capacity after decidualization. Under replication stress, expression of all three MRN complex components is reduced in Endo MenSCs, a finding also observed in scRNA-seq data from endometriosis patient tissue. Consequently, Endo MenSCs accumulate more regions of under-replicated DNA and unrepaired damage, promoting senescence and reducing clonogenicity. Thus, in some patients with endometriosis, eutopic MenSCs experience varying levels of replication stress and exhibit an impaired ability to mount an effective DDR; thereby leaving a subset of cells with proliferative potential that could contribute to the development of ectopic endometrial lesions. Altogether, this study raises the possibility that the defective DDR in endometriosis patients can be exploited for the identification of a non-hormonal therapeutic that reduces the growth of endometriotic cells.

## Author’s Roles

K.C., R.M., J.M., L.H., developed the concepts and supervised the research; K.C., J.M., R.M. designed experiments; K.C., G.T. performed the experiments; K.C., G.T., H.E.S. analyzed data; L.H. gave technical support and conceptual advice; K.C., L.H. critically discussed the data; L.H. acquired funding; K.C., L.H. wrote the original manuscript; K.C., L.H., G.T., J.M. R.M., H.E.S. reviewed and edited the manuscript. All authors have read and agreed to the published version of the manuscript.

## Acknowledgements

We acknowledge core support from the Institute for the Biology of Stem Cells at University of California, Santa Cruz and California Institute for Regenerative Medicine (CIRM) Shared Stem Cell Labs (RRID:SCR_021353), Microscopy (RRID:SCR_021135), Facilities (CL1-00506-1,2), and Major Facilities (FA1-00617-1) awards. We acknowledge core support from the University of California, Santa Cruz Chemical Screening Center (RRID:SCR_021114), and support from the National Institutes of Health (1S10RR022455-01A1, 1S10OD028730-01A1). We also acknowledge core support from University of California, San Francisco grant (1S10OD036282-01) and University of California, Berkeley QB3 High Throughput Screening Core Facility (RRID:SCR_022304). This work was supported by the National Institutes of Health: Eunice Kennedy Shriver National Institute of Child Health and Human Development (NIHCD) under award number R01HD098722 to L.H. and T32HD108079 to J.M. Other support for individuals was from a California Institute of Regenerative Medicine (CIRM) training grant (EDUC-12759) to R.M and from the UCSC Koret Scholarship (K.C. and G.T.). We thank Ben Abrams, Bari Nazario, Patty Lovelace, Beverley Rabbitts, and Mary West for technical assistance. We thank Santa Cruz Biotechnology for antibodies.

## Figure Legends>

**Supplementary Figure 1: A)** Representative flow cytometry plot showing selection of the MenSCs population. **B-F**) Representative flow cytometry plot and percentage of positive cells for CD140b (**B**, **C**), SUSD2 (**D, E)** and CD146 (**F**). Light gray dots are FMO negative controls used to make positive gates (black boxes) and dark gray dots are full-stained cells. **G**) Representative flow cytometry histogram showing gates for G1, S, and G2/M phases of the cell cycle. **H**) Percentage of proliferating healthy and Endo cells in G1, S, or G2/M phases of the cell cycle, as detected by flow cytometry analysis. **I**) Percentage of proliferating healthy and Endo cells positive for Ki67, as detected by ICC. **J**) Representative flow cytometry histogram showing the gate for EdU positive cells. The light gray peak represents the DMSO negative control used to make the positive gate. The dark gray peaks represent cells incubated with EdU. **K**) Percentage of proliferating healthy and Endo cells positive for EdU, as detected by flow cytometry analysis. **L**) Quantification of nuclear γH2AX integrated intensity in UnDec healthy and Endo sample, as detected by ICC. For (**I**), data represents the means of *n*=3 biologically independent samples quantified from 9 fields of view (∼1,000 nuclei). For (**L**), black dots represent the means of *n*=3 biologically independent samples and gray dots represent the individual nuclei of each sample quantified from 9 fields of view (∼1,000 nuclei). Data representative of *n*=3 biologically independent samples (**C**, **E**, **H**, **K**). Data presented as mean ± SD (**C**, **E**, **H**, **I**, **K**) and mean ± SEM (**L**). Data analyzed by two-way ANOVA with Tukey’s (**C, E**) or Šídák’s (**H**) multiple comparison test and Welch’s, two-tailed *t* test (**I**, **K**, **L**). *p* value: ns>0.05.

**Supplementary Figure 2: A**) Quantification of nuclear γH2AX integrated intensity in UnDec healthy and Endo samples, as detected by ICC. **B**) Percentage of nuclei with focal or diffuse nuclear γH2AX in UnDec healthy and Endo samples. **C**) Quantification of nuclear pATM integrated intensity in UnDec healthy and Endo samples, as detected by ICC. **D**) Percentage of UnDec healthy and Endo cells in G1, S, or G2/M phases of the cell cycle, as detected by flow cytometry analysis. **E**) Quantification of nuclear 53BP1 integrated intensity in UnDec healthy and Endo samples, as detected by ICC. **F**) Quantification of the number of 53BP1-NB foci per focal nucleus in UnDec healthy and Endo samples. For (**A, C, E, F**), black dots represent the means of *n*=3 biologically independent samples and gray dots represent the individual nuclei of each sample quantified from 9 fields of view (∼1,000 nuclei) or 27 fields of view (∼500 nuclei). For (**B**), bars represent the means of *n*=3 biologically independent samples quantified from 27 fields of view (∼500 nuclei). Data representative of *n*=3 biologically independent samples (**D**). Data presented as mean ± SD (**B, D**) and mean ± SEM (**A**, **C**, **E**, **F**). Data analyzed by two-way ANOVA with Šídák’s multiple comparison test (**D**) and Welch’s, two-tailed *t* test (**A, B, C, E, F**). *p* value: ns>0.05.

**Supplementary Figure 3: A**) MTT assay of UnDec healthy (red, circles) and Endo (black, squares) cells after Hu-treatments showing no differences between healthy and Endo samples, presented as optical density. **B**) MTT assay of UnDec Endo cells after Hu-treatments showing concentration that affects cell viability (30mM), presented as optical density. **C**) Percentage of nuclei with focal or diffuse nuclear γH2AX in Hu-treated, UnDec healthy and Endo samples. **D**) Quantification of nuclear pATM integrated intensity in untreated and Hu-treated, UnDec healthy and Endo samples, as detected by ICC. **E**) Representative images of pATM (white) from untreated (top) and Hu-treated (bottom), UnDec healthy (left) and Endo (right). DNA by Hoechst in blue. **F**) Percentage of nuclei with focal or diffuse nuclear pATM in Hu-treated, UnDec healthy and Endo samples. **G**) Quantification of the number of pATM foci per focal nucleus in Hu-treated, UnDec healthy and Endo samples. **H**) Quantification of pATM foci area in Hu-treated, UnDec healthy and Endo samples. **I**) Quantification of nuclear pCHK2 integrated intensity in untreated and Hu-treated, UnDec healthy and Endo sample, as detected by ICC. **J**) Representative images of pCHK2 (white) from untreated (top) and Hu-treated (bottom), UnDec healthy (left) and Endo (right) samples. DNA by Hoechst in blue. **K**) *CDKN1A* (*p21*) gene expression in untreated and Hu-treated, UnDec healthy and Endo samples, as detected by RT-qPCR. **L**) Quantification of nuclear p21 integrated intensity in untreated and Hu-treated, UnDec healthy and Endo sample, as detected by ICC. **M**) Representative images of p21 (white) from untreated (top) and Hu-treated (bottom), UnDec healthy (left) and Endo (right) samples. DNA by Hoechst in blue. For (**C, F**), bars represent the means of *n*=3 biologically independent samples quantified from 27 fields of view (∼150 nuclei). For (**G**), black dots represent the means of *n*=3 biologically independent samples and gray dots represent the individual nuclei of each sample quantified from 27 fields of view (∼150 nuclei). Data presented as mean ± SD (**A**, **B**, **C**, **F**, **K**) and mean ± SEM (**D**, **G**, **H**, **I**, **L**). Data representative of *n*=3 biologically independent samples (**A**, **B**, **D**, **H**, **I**, **K**, **L**). Data analyzed by ordinary one-way ANOVA with Tukey’s multiple comparison test (**B**), two-way ANOVA with Tukey’s (**D**, **I**, **K**, **L**) or Šídák’s (**A**) multiple comparison test and Welch’s, two-tailed *t* test (**C**, **F**, **G**, **H**). *p* values: ns>0.05, **<0.01, ***<0.001, ****<0.0001.

**Supplementary Figure 4: A**) Representative images of untreated (top) and Hu-treated (bottom), Dec healthy (left) and Endo (right) cells. Vimentin detected in magenta and DNA by Hoechst in blue. **B**) Representative images of SA-β-Gal staining (blue) from untreated (top) and Hu-treated (bottom), Dec healthy (left) and Endo (right) samples.

