## Supplementary figures and images for "Defects in the DNA Damage Response of Patient-derived Endometriosis Stromal Cells"

### Supplemental Figures 1-4

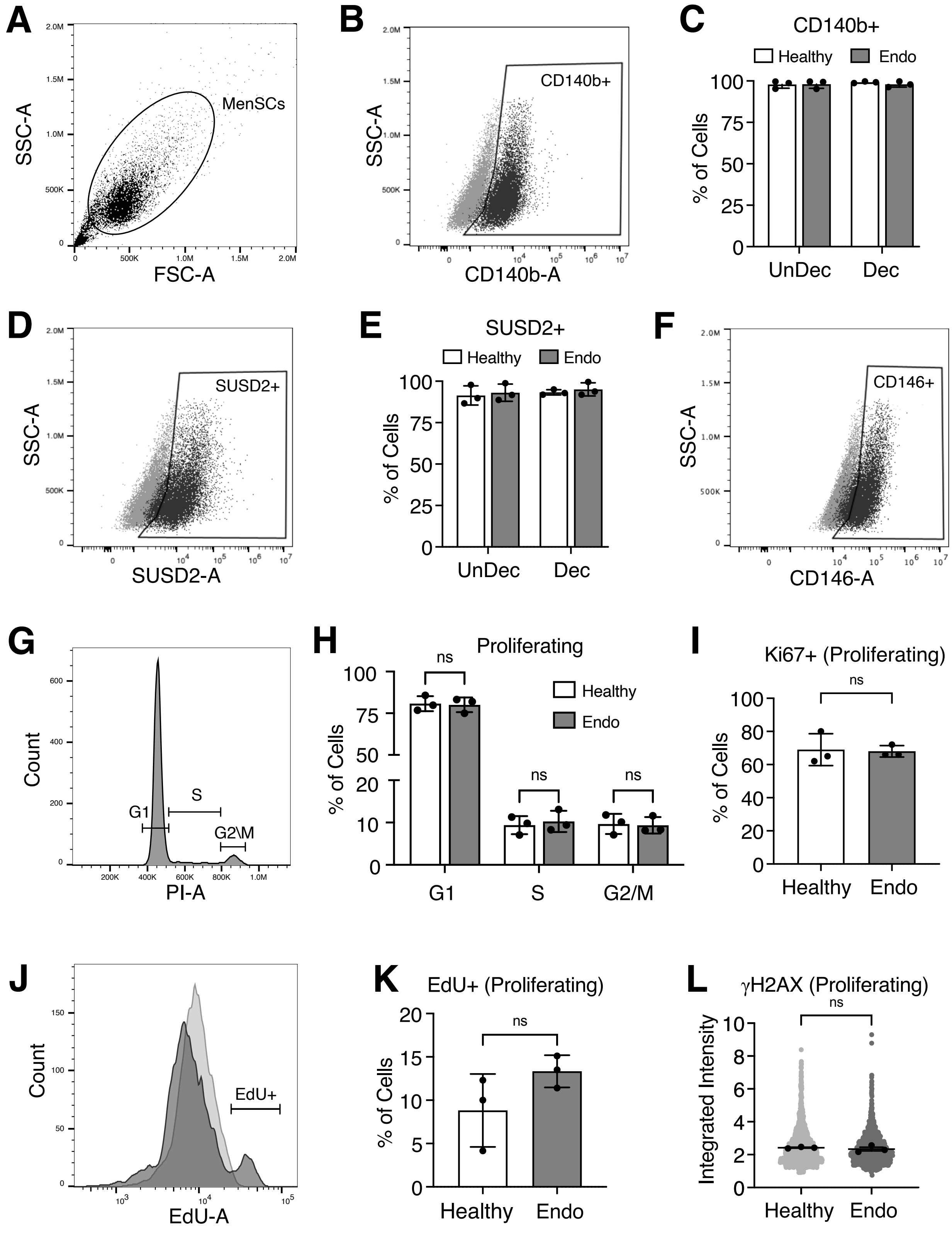


Suppl. Figure 1

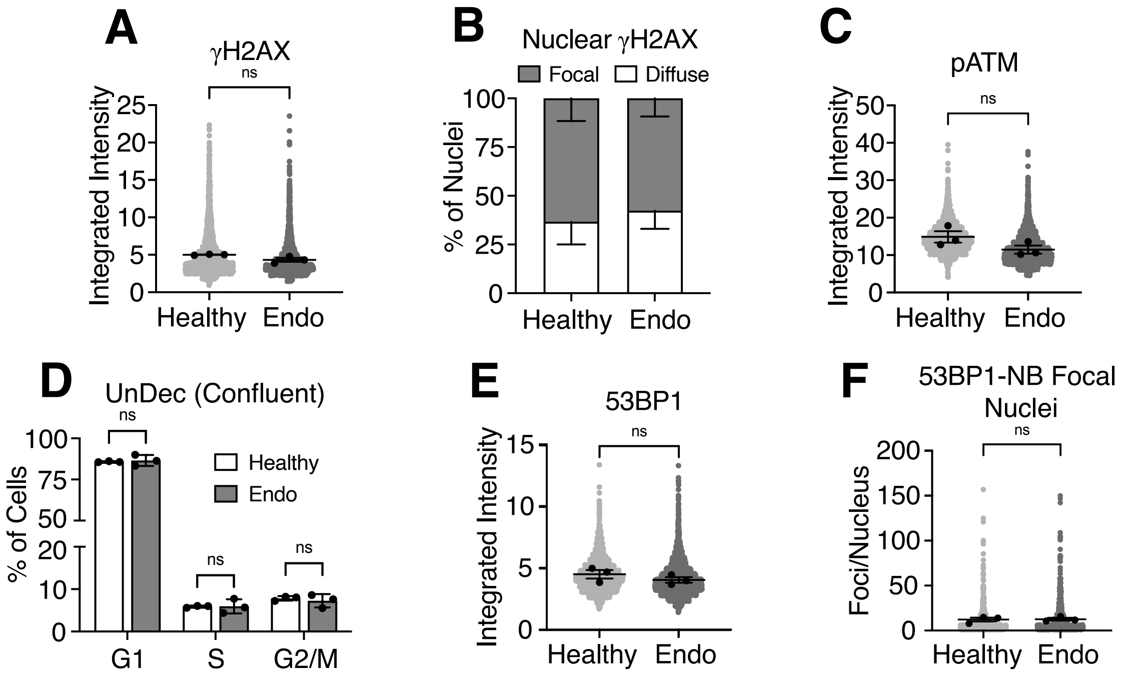


Suppl. Figure 2

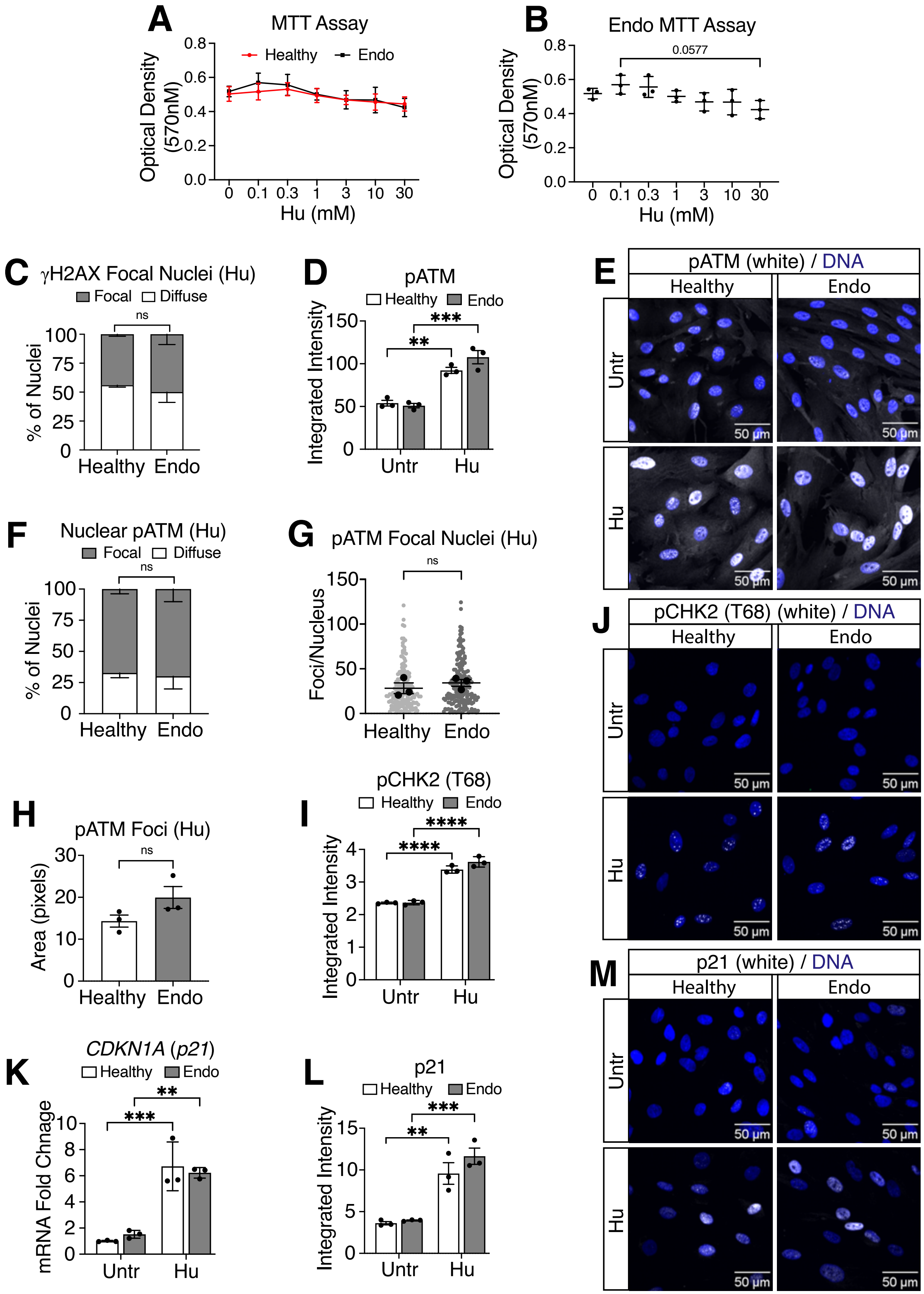


#### Suppl. Figure 3

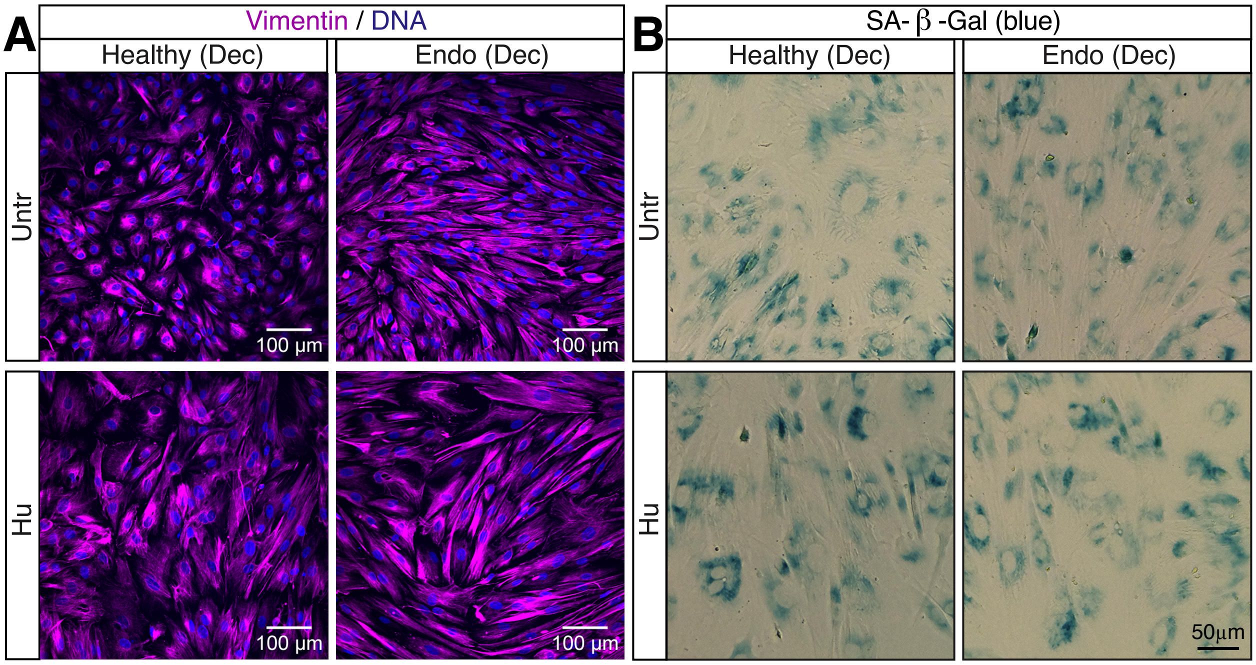


Supp Figure 4
