## Supplementary material for "Defects in the DNA Damage Response of Patient-derived Endometriosis Stromal Cells": Table 1

Table 1: Study participant descriptions and healthy histories.

| | Sample 1 | Sample 2 | Sample 3 | Sample 4 | Sample 5 | Sample 6 | Sample 7 |
|---|---|---|---|---|---|---|---|
| Age | 34 | 21 | 36 | 37 | 45 | 42 | 39 |
| Length of menstrual cycle | 20 days | 27 days | N/A | 25 days | N/A | N/A | 28-35 days |
| Hormonal birth control | No | No | No | No | No | No | No |
| Diagnosed endometriosis? Category? | No | No | No | No | Yes; 4 | Yes; 1 | Yes, N/A |
| Family history of endometriosis? | No | No | No | No | No | No | No |
| Diagnosed PCOS? | No | No | No | No | No | No | Yes |
| Family history of PCOS? | No | No | No | No | No | Yes | No |
| Diagnosed endometrial cancer? | No | No | No | No | No | No | No |
| Family history of endometrial cancer? | No | No | No | No | No | No | No |
| Diagnosed ovarian cancer? | No | No | No | No | No | No | No |
| Family history of ovarian cancer? | No | No | Yes | No | No | No | No |
| Full-term pregnancy? | No | No | Yes | Yes | No | Yes | Yes |
| Miscarriages? | No | No | Yes | Yes | No | No | Yes |
